# Xeno-free human iPSC-derived prostate organoid platform for multilineage differentiation and genetic manipulation

**DOI:** 10.1101/2025.10.21.683654

**Authors:** Neha Shaikh, Matthew Teasdale, Lewis J Walker, Laura Wilson, Rachel Howarth, Sara Saleem, Anastasia C Hepburn, Ryan Nelson, Qiuyu Lian, Rafiqul Hussain, Jonathan Coxhead, Luke Gaughan, Majlinda Lako, Emma Scott, Craig Robson, Ben Simons, Simon W. Hayward, Douglas W. Strand, Rakesh Heer, Adriana Buskin

## Abstract

Current prostate organoid models rely on tissue-derived material or animal components and lack epithelial and stromal complexity. We defined a xeno-free system to generate human prostate organoids from induced pluripotent stem cells with consistent multilineage differentiation. Floating organoids self-organize into epithelial and stromal domains with basal, luminal, neuroendocrine, fibroblast, and smooth muscle markers. In an alternative modular co-culture system, engineered epithelial progenitors are aggregated with wild-type mesenchymal progenitors, enabling compartment-specific manipulation. Androgen receptor–overexpressing organoids showed increased epithelial AR and PSA expression and proliferation. Single-cell transcriptomics, together with qPCR and immunostaining, confirmed prostate lineage specification and tissue organization. This new xeno-free platform provides a reproducible, scalable, and genetically tractable model to study in-vitro prostate lineage programs, epithelial-stromal interactions, and disease biology.

**Graphic Abstract:** This study describes the generation of prostate organoids from human iPSCs. iPSCs, including those reprogrammed from patients carrying germline mutations, can be differentiated into prostate organoids either through monoculture or by co-culturing endodermal cells with mesenchymal progenitors. Genetic manipulation can be introduced before endoderm specification to model cancer drivers. The resulting multi-lineage organoids exhibit distinct epithelial (AR⁺, NKX3.1⁺, PSA⁺, CK8/18⁺) and stromal (VIM⁺, α-SMA⁺) compartments, providing a versatile platform for developmental studies, disease modelling, drug screening, and biomarker discovery.

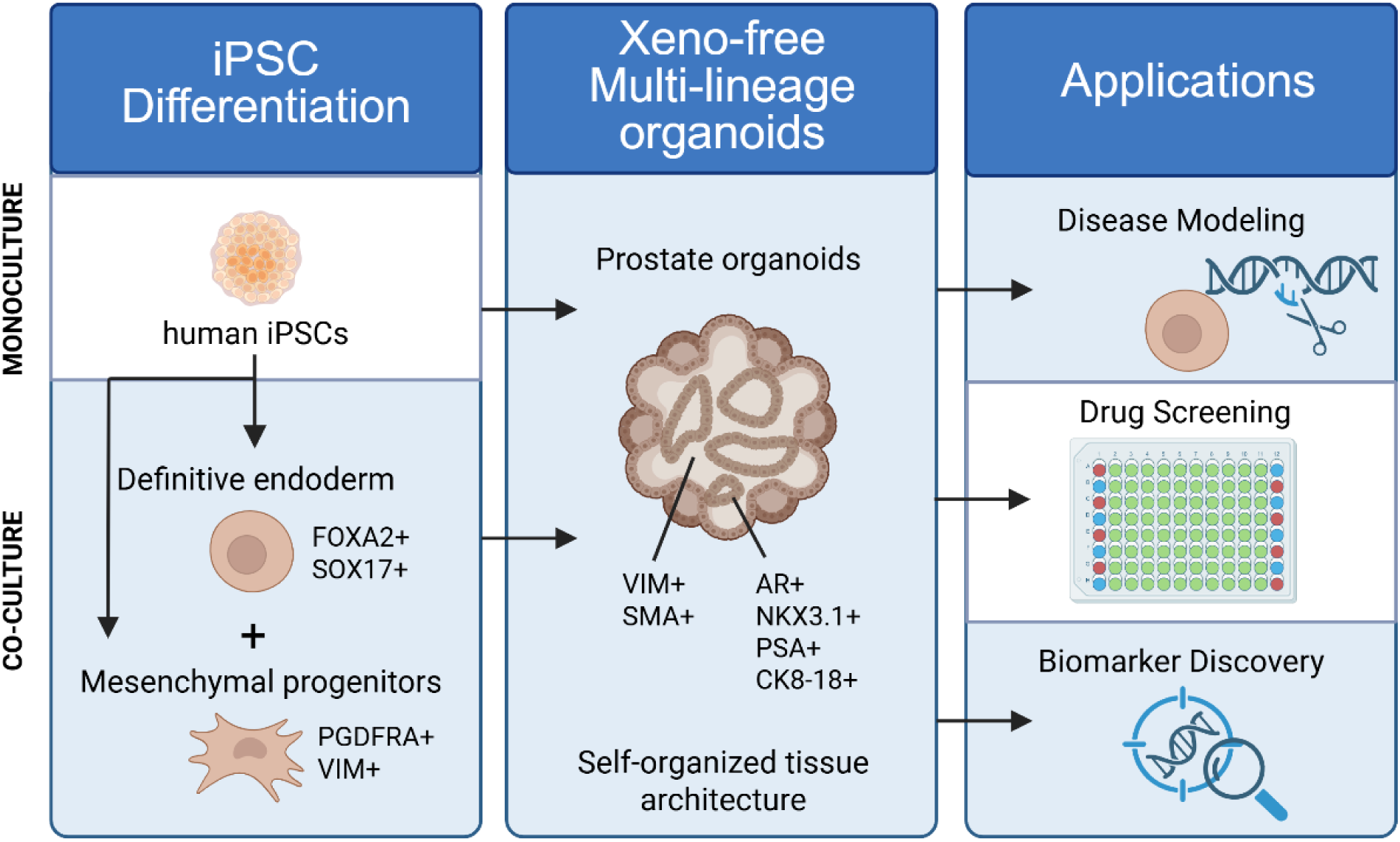

## Introduction

Prostate diseases affect hundreds of millions of men globally, with benign prostatic hyperplasia (BPH), prostate cancer, prostatitis, and lower urinary tract symptoms imposing a substantial and growing health burden (1,2). Although an area of intense research, progress is limited by the scarcity of physiologically relevant, human-specific models that accurately reflect the complexity of tissue histology and cellular differentiation. For example, prostate cancer researchers rely on fewer than ten well-characterized prostate cancer cell lines. While these lines have contributed substantially to our understanding of androgen receptor (AR) signal targeting as a central approach to treating advanced prostate cancer, they are inherently limited by their two-dimensional structure and the absence of tumor heterogeneity and stromal–epithelial interactions (3).

Tumor slice cultures offer the advantage of preserving native tissue architecture, but they are short-lived, difficult to scale, and exhibit transcriptomic stress responses that cloud experimental analysis (4). Genetically engineered mouse models (GEMMs) and patient-derived xenografts (PDXs) remain invaluable for *in vivo* validation, yet both differ from human prostate tissue in stromal composition, AR signaling, and treatment response (5–8). GEMMs have distinct mouse biology that is different from human. PDX models, in contrast, retain some human biology although stromal environment changes with increasing passage number, and grafts require either immunodeficient or humanized hosts with no or limited immune responses. Animal-based models face constraints related to cost, engraftment efficiency, and long-term stability (9,10)

Patient-derived organoids (PDOs) have significantly advanced the field by preserving patient-specific genomic and phenotypic features (11–13). However, PDO systems remain limited by their reliance on advanced-stage or metastatic samples, inefficiency in culturing treatment-naïve tumors, and poor preservation of the stromal compartment (14–16). While 3D organoid technology has transformed disease modeling in other epithelial tissues, robust, isogenic, scalable, and experimentally tractable models to guide personalized therapy in prostate cancer are still lacking (17). Recent studies have begun to address these limitations by incorporating stromal components into culture systems and by generating human organoids from the differentiation of pluripotent stem cells as an alternative to tissue-derived material (18,19).

Tissue recombination studies have shown that rodent urogenital mesenchyme (UGM) can direct the differentiation of human prostate epithelial cells into organized, functional prostate tissue *in vivo*. These models demonstrated that stromal–epithelial interactions are essential for prostate organogenesis, supporting the development of basal and luminal epithelial lineages within a mesenchymal-derived stromal compartment formed by the inductive rodent UGM (20,21). Building on these principles, we previously demonstrated that human induced pluripotent stem cells (iPSCs) can be directed toward prostate epithelial fate through co-culture with rat UGM, generating prostate tissue both *in vivo* and *in vitro* that included neuroendocrine cells and a supporting stromal niche, thereby reproducing multiple cell types found in human prostate tissue (22). While this established the developmental potential of iPSCs for prostate modeling, the use of animal-derived components such as UGM and Matrigel limit reproducibility, scalability, affordability and translational applicability.

In the present study, we present a fully defined, xeno-free protocol for generating human prostate organoids from induced pluripotent stem cells. These animal-free approaches yield self-organizing structures that capture key aspects of prostate lineage specification and tissue organization, with epithelial and stromal compartments arranged in distinct domains. The platform is reproducible and scalable, and supports studies of differentiation, stromal–epithelial interaction, lineage specification, and disease modeling.

## Results

### Prostate Organoids Self-organize and Recapitulate Prostate Architecture

We generated three iPSC lines from healthy prostate tissues obtained from cystoprostatectomy surgery for bladder cancer as previously described (22). All three cell lines were confirmed to be iPSCs by confirming pluripotency and trilineage differentiation potential (Figure S1). To initiate prostate differentiation, iPSCs were aggregated into spheroids in low attachment 96-well plates. On day 1, 3D spheroids were treated with Activin A, to induce endodermal differentiation. By day 3, they expressed *SOX17* and *T (Brachyury)*, confirming an early mesendodermal state (Supplementary Figure 2A). At this point, spheroids were switched to prostate specification medium for 4 days, followed by prostate organoid medium for the remainder of the differentiation protocol (Figure 1A).

**Figure 1.**
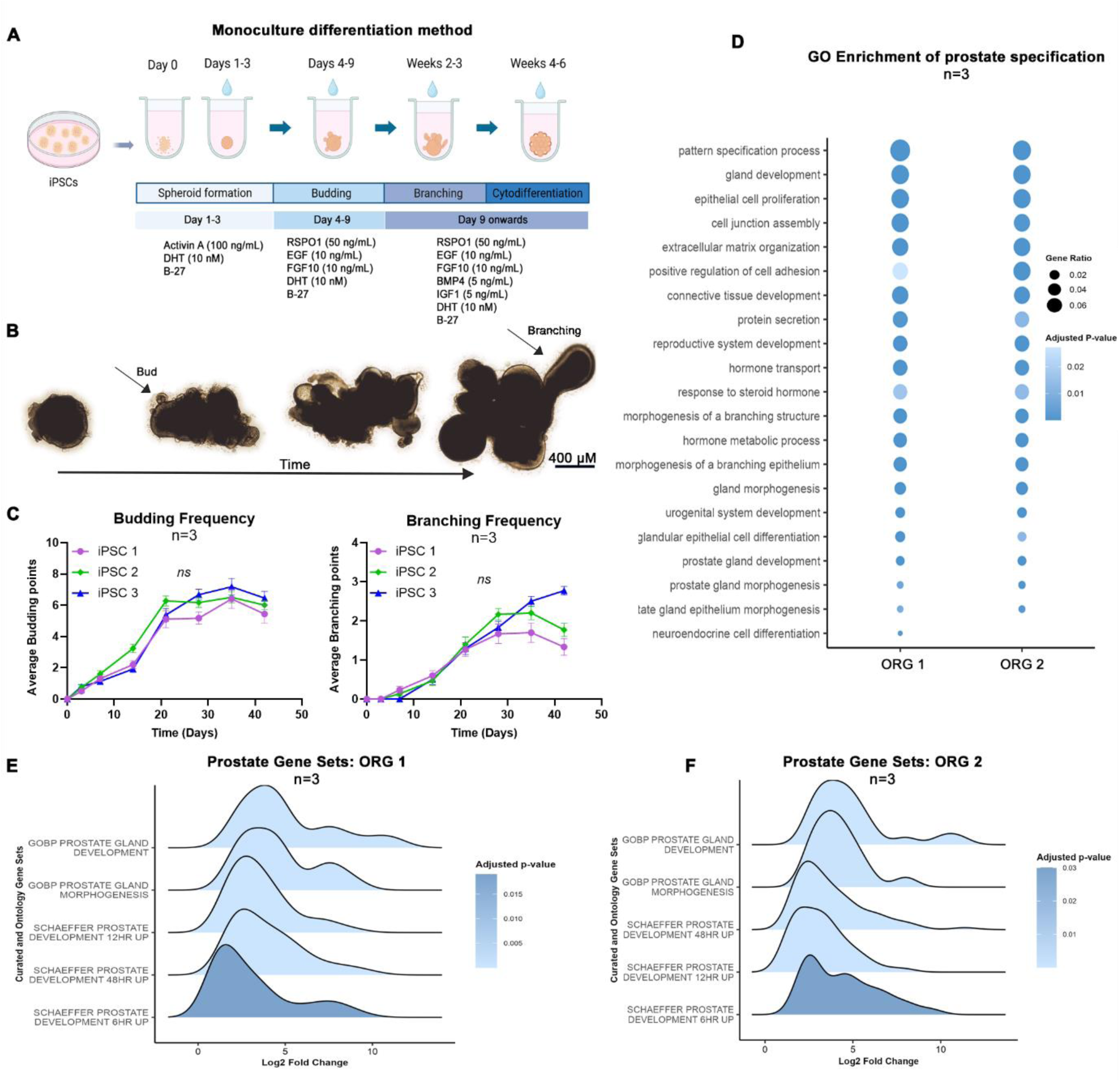
Human iPSC-derived prostate organoids self-organize and recapitulate key features of prostate development at week 6. (A) Schematic of the stepwise differentiation protocol from iPSCs, illustrating key stages of spheroid formation, budding, branching, and cytodifferentiation. (B) Representative brightfield images showing progressive organoid growth and morphogenesis from week 1 to week 6. Scale bar, 400 µm. (C) Quantification of budding and branching frequencies (n=3) across three independent iPSC lines at week 6. Budding refers to the emergence of a single protrusion from the organoid surface, while branching involves further extension or bifurcation of these protrusions. A total of 90 organoids were assessed (30 per iPSC line: iPSC1, iPSC2, and iPSC3). These iPSC lines were derived from different donors and used as biological replicates to assess the consistency of developmental outcomes. (D) Gene Ontology (GO) enrichment analysis (n=3) of bulk RNA-seq data from week 6 organoids derived from iPSC1 and iPSC2 reveals consistent biological processes related to prostate development. (E–F) Gene Set Enrichment Analysis (GSEA) (n=3) of week 6 organoids derived from iPSC1 (E) and iPSC2 (F) versus their undifferentiated iPSC counterparts shows significant upregulation of prostate development gene signatures. The top-enriched gene sets include human Gene Ontology terms related to prostate gland development and morphogenesis. Additional enrichment of Schaeffer prostate development signatures, originally defined in a mouse model, highlights conserved androgen-responsive transcriptional programs recapitulated in the human organoids.

Throughout the 6-week differentiation period, organoids were live-imaged to monitor morphological changes. Quantification of budding and branching events at week 6 (Figure 1B-C) revealed consistent developmental outcomes across all three iPSC lines (n=3; 30 organoids per iPSC line), demonstrating high reproducibility. No significant differences were observed among lines.

Two iPSCs and their derived organoids were characterized via transcriptomic analyses at week 6 and revealed a clear transcriptional divergence between undifferentiated iPSCs and differentiating prostate organoids using principal component analysis (Supplementary Figure 3A). Gene Ontology analysis (Figure 1D) showed enrichment of biological processes associated with prostate specification. Gene Set Enrichment Analysis (Figure 1E-F) further revealed strong upregulation of gene signatures linked to prostate gland differentiation. Notably, enrichment of the Schaeffer androgen responsive prostate development signature, originally defined in mouse, highlights the conserved nature of these developmental programs in our human organoids (23).

### Acquisition of Epithelial and Stromal Prostate Lineages

Organoids at week 6 were fixed, processed, and paraffin embedded for histological analysis. Immunohistochemistry revealed distinct epithelial and stromal compartmentalization within the organoids. Epithelial nests expressed key prostate lineage-specific markers NKX3-1 and PSA, and confirmed the presence of luminal cells expressing CK8/18 and AR, neuroendocrine cells expressing chromogranin A, as well as a basal epithelial layer expressing p63. These epithelial structures were surrounded by a stromal compartment positive for vimentin and α-SMA, consistent with normal human prostate histology (24,25) (Figure 2A-B).

**Figure 2.**
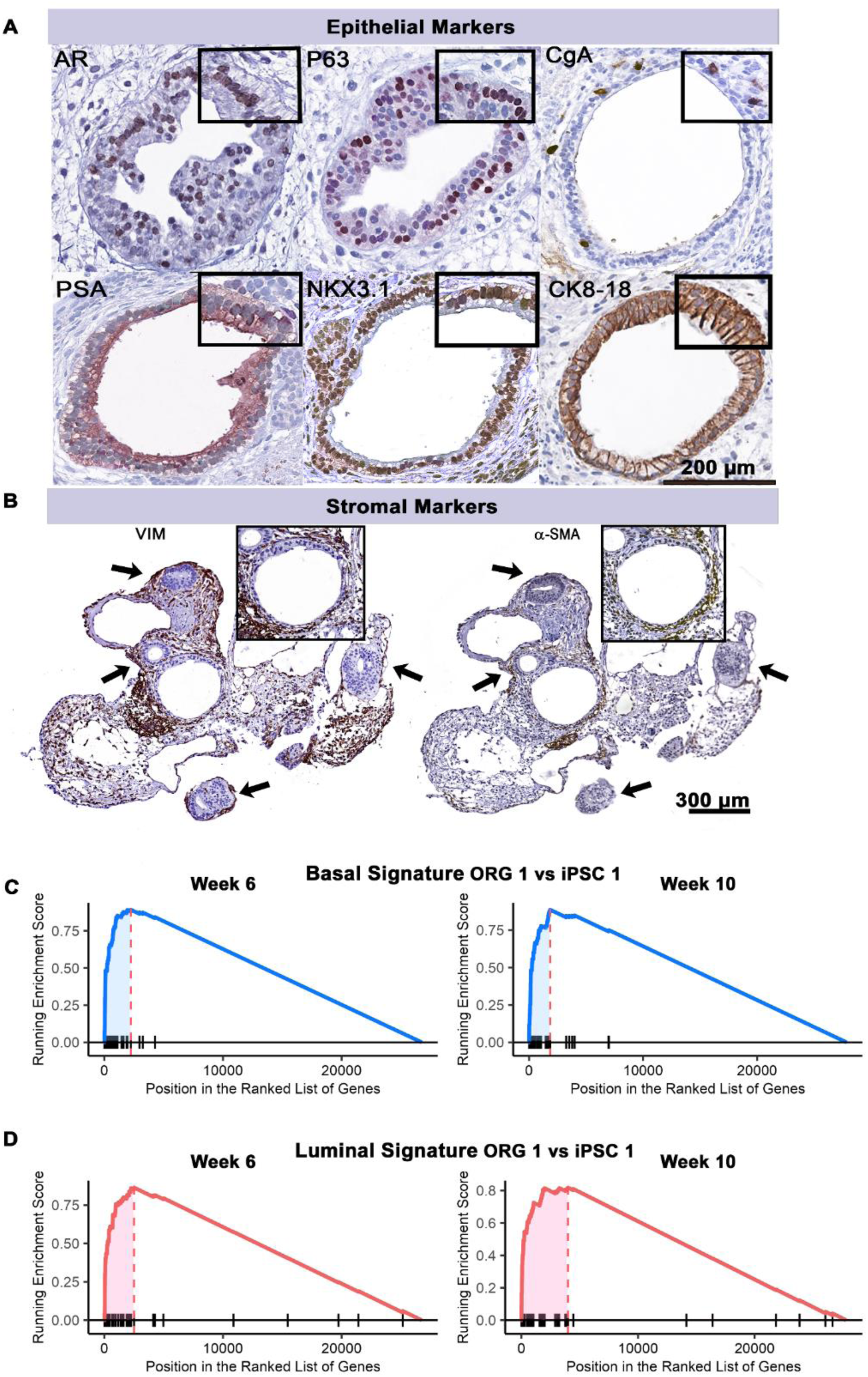
Prostate organoids recapitulate epithelial and stromal compartmentalization with progressive acquisition of tissue-specific gene signatures. (A–B) Immunohistochemical staining of whole organoid sections reveals distinct epithelial regions expressing androgen receptor (AR), basal marker P63, neuroendocrine marker chromogranin A (CgA), prostate-specific antigen (PSA), NKX3-1, and cytokeratins 8 and 18 (CK8/18), supporting basal, luminal, and neuroendocrine differentiation. Immunostaining for stromal markers shows positive staining for vimentin (VIM) and α-smooth muscle actin (α-SMA). Black arrows indicate epithelial nests surrounded by stromal compartments. (C) Gene set enrichment analysis (GSEA) using basal signatures derived from primary adult prostate epithelial cells in organoids (ORG1) versus undifferentiated iPSCs (iPSC1) at weeks 6 and 10, showing enrichment of basal genes. (D) GSEA using luminal signatures derived from primary adult prostate epithelial cells (n = 3) at weeks 6 and 10 shows progressive enrichment, consistent with luminal lineage specification over time.

Transcriptomic analysis revealed epithelial lineages consistent with prostate lineage specification. Basal signatures included *TP63* and *KRT5*, while luminal signatures encompassed *FOXA1*, *AGR2*, *CEACAM6*, *TFF3*, and *KRT19*. Neuroendocrine-associated transcripts such as *CHGA*, *SYP*, and *ENO2* were also detected (Supplementary Figure 3C-D). Classical epithelial keratins CK8 (*KRT8*) and CK18 (*KRT18*) were abundantly expressed and readily detected by immunohistochemistry, although their transcript levels varied between clones, with evidence of increased expression in ORG2 organoids. Gene set enrichment analysis (GSEA) using basal and luminal signatures derived from primary adult prostate epithelial cells further supported these lineage identities, showing significant enrichment of basal epithelial programs at week 6 and enhanced acquisition of luminal signatures at week 10 (Figure 2 C-D). This temporal pattern indicates a transition from basal-enriched to luminal-enriched states, mirroring the maturation trajectory described by Hepburn *et al.* (22), where iPSC-derived prostate organoids advance along epithelial differentiation pathways during development.

Beyond the epithelial lineage, stromal gene expression, including *VIM*, *FAP*, *ACTA2*, *PDGFRB*, and *PDGFRA*, was maintained or increased during maturation, consistent with the histological identification of a well-developed stromal compartment surrounding epithelial nests. Notably, we observed robust upregulation of extracellular matrix genes such as *LAMB1*, *ECM1*, *ECM2*, *COL22A1*, and *COL6A6*, highlighting the capacity of these organoids to self-organize and synthesize their own extracellular matrix under defined scaffold-free conditions (26,27). Together, these profiles reflect a multicompartmental architecture with epithelial, stromal, and extracellular-matrix-producing elements in defined domains (Supplementary Figure 3B-D).

Originating from a common iPSC progenitor, this monoculture system is well suited for developmental studies and for modeling inherited conditions. It also enables the introduction of germline cancer-risk alleles, ensuring that both epithelial and stromal compartments share the same genetic context.

### Androgen Supplementation is Essential for Epithelial Differentiation

To assess the contribution of androgen signaling to prostate organoid differentiation, we cultured organoids with or without dihydrotestosterone (DHT) and assessed marker expression at weeks 4 and 6 using immunohistochemistry.

Bar plots were generated from IHC images (n=3) to quantify the marker-positive area. This analysis showed that organoids cultured with DHT exhibited increased expression of epithelial markers CK8/18 and p63 over time, while those in androgen-deprived conditions showed reduced expression of both markers (Figure 3A-B). In contrast, expression of stromal markers α-SMA and vimentin was maintained or elevated in the absence of DHT, indicating that stromal differentiation does not depend on androgen signaling (Figure 3C-D).

**Figure 3.**
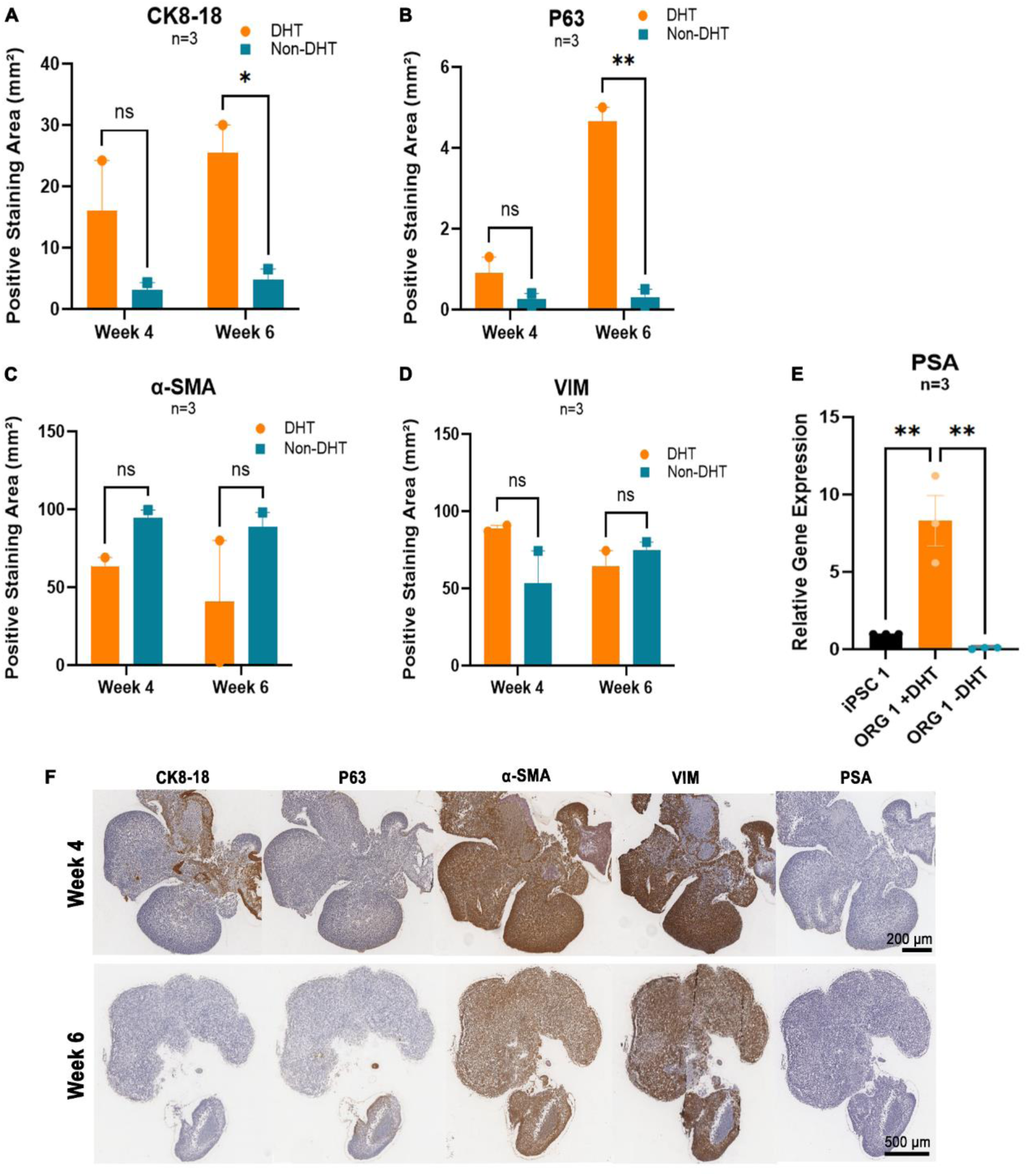
Androgen supplementation enhances epithelial differentiation in iPSC-derived prostate organoids. (A–B) Bar plots showing positive staining area for epithelial markers CK8/18 (A) and P63 (B) at weeks 4 and 6. DHT-treated organoids (orange bars) show increased staining over time, while androgen-deprived conditions (blue bars) show reduced epithelial marker expression. (C–D) Equivalent plots for stromal markers α-SMA (C) and VIM (D) indicate that stromal marker expression is preserved or increased in the absence of DHT. (E) Quantification of *KLK3* (PSA) transcript expression by qPCR (n = 3) shows reduced levels in androgen-deprived conditions compared with DHT-treated organoids. iPSC1 is shown as a baseline control. Data are presented as mean ± SEM; *p < 0.05. (F) Representative immunohistochemistry images of whole organoid sections at weeks 4 and 6 cultured without DHT, stained for epithelial markers CK8/18, P63, and PSA, and stromal markers á-SMA and VIM. Control images at week 6 are shown in Figure 2A-B.

Quantification of *KLK3* (PSA) expression by qPCR (n = 3) showed a significant reduction under androgen-deprived conditions compared to DHT-treated organoids, with iPSC1 included as a baseline reference (Figure 3E; mean ± standard error of the mean (SEM), *p < 0.05). Representative IHC images confirmed these observations. Organoids cultured without DHT displayed markedly lower staining for CK8/18, p63, and PSA, while α-SMA and vimentin expression remained detectable, consistent with previous reports that stromal markers are less affected by androgen withdrawal (28–30).

### Co-culture Strategy Enables Compartment-specific Manipulation

The above model design involves the differentiation of prostate organoids and their epithelial and stromal tissues from a common progenitor. To establish utility for disease modeling, such as a prostate cancer organoid in which mutations would need to be confined to the epithelium as seen in patients, we developed an alternative differentiation protocol. Here, we tested whether we could “assemble” prostate organoids by combining definitive endodermal progenitors (epithelial precursor) and mesenchymal progenitors (stromal precursors) for organoid differentiation, enabling independent genetic manipulation of each compartment. This approach involved the directed differentiation of iPSCs into lineage-specific progenitors, followed by aggregation into spheroids and subsequent specification into prostate organoids (Figure 4A).

**Figure 4.**
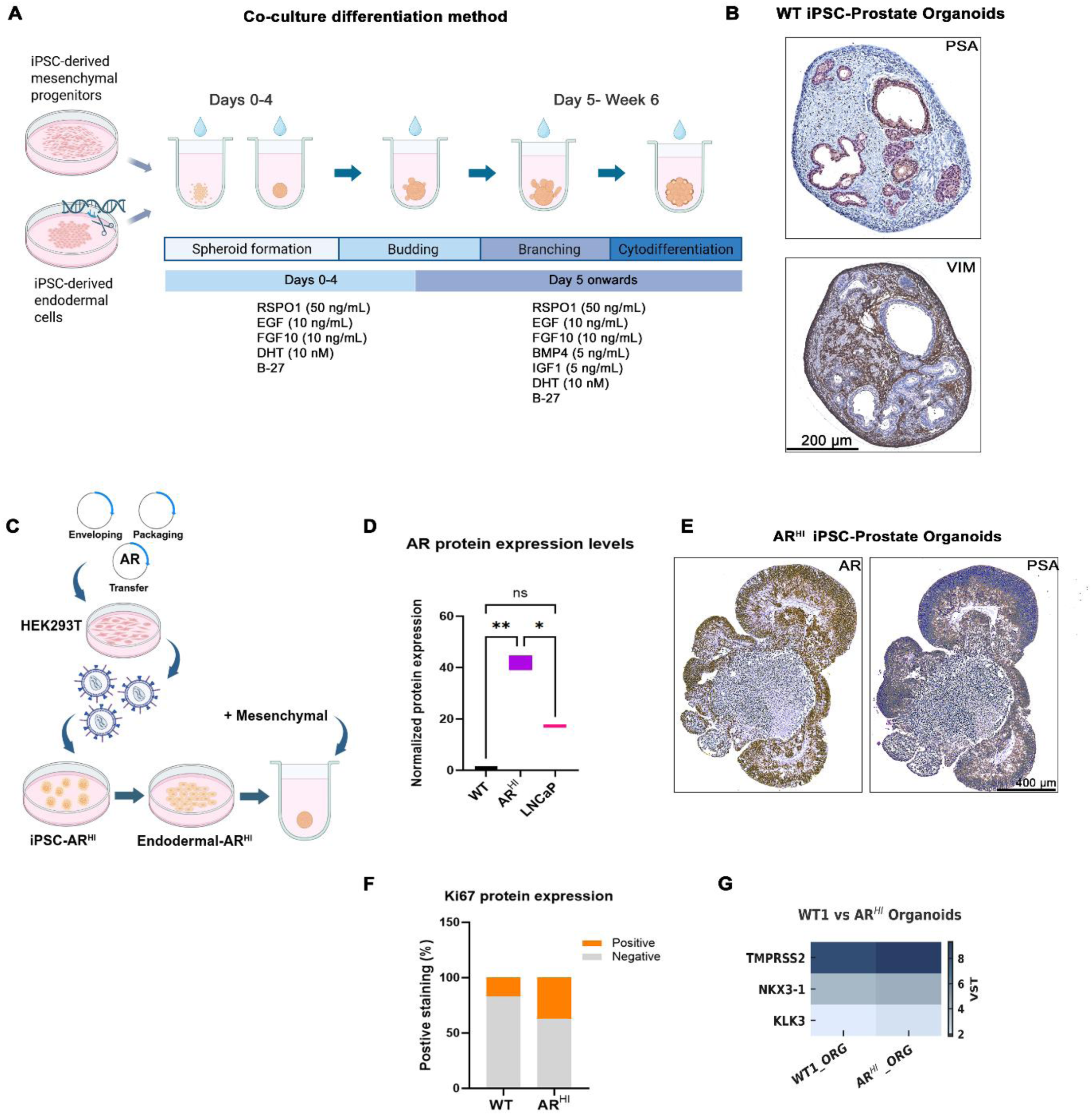
Co-culture model combining engineered endodermal cells and mesenchymal progenitors enables prostate organoid generation and epithelial compartment-specific genetic manipulation. (A) Schematic of the co-culture protocol using iPSC-derived mesenchymal progenitors and engineered iPSCs differentiated into endodermal cells. This system enables spatial compartmentalization of stromal and epithelial lineages, supporting targeted manipulation of the epithelial compartment. Developmental stages are shown: spheroid formation, budding, branching, and cytodifferentiation. (B) Immunohistochemistry of WT organoids demonstrates PSA-positive epithelial regions and VIM-positive stromal regions. (C) Overview of the epithelial-specific manipulation strategy. The *AR* transgene was introduced into iPSCs, which were differentiated into endodermal cells (Endodermal-*AR*ᴴᴵ), and co-cultured with mesenchymal progenitors to generate prostate organoids. (D) Quantification of AR protein levels by western blotting in iPSC-WT, iPSC-*AR*ᴴᴵ, and LNCaP cells, showing increased AR expression in iPSC-*AR*ᴴᴵ compared with WT controls. (E) Immunohistochemistry of *AR*ᴴᴵ organoids shows epithelial expression of AR and PSA, confirming epithelial targeting. (F) Ki67 staining reveals a higher proportion of proliferating cells in *AR*ᴴᴵ organoids compared with WT controls. (G) Heatmap of variance-stabilized (VST) expression values shows induction of canonical AR targets (*TMPRSS2*, *NKX3-1*, *KLK3*) in *AR*ᴴᴵ organoids.

At day 3, gene expression analysis confirmed successful induction of definitive endoderm, as shown by increased expression of *SOX17* and *FOXA2* in 2D iPSC cultures. Downregulation of pluripotency markers *NANOG* and *OCT4* confirmed loss of iPSC identity during differentiation. By day 21, mesenchymal progenitors showed high expression of *NCAM1* and reduced levels of epithelial (*EpCAM*) and pluripotency markers (*NANOG*, *OCT4*). *T (Brachyury)* and *MIXL1*, transient markers associated with early mesoderm and mesendoderm specification, were also downregulated, indicating progression towards a committed mesenchymal fate (Supplementary Figure 2). Immunohistochemistry of co-cultured organoids at week 10 revealed clear compartmentalization, with PSA-positive epithelial regions and vimentin-positive stromal regions (Figure 4B).

As proof of concept for epithelial-specific genetic manipulation, we introduced an AR transgene into iPSCs prior to endoderm differentiation. These engineered cells (Endodermal-AR^HI^) were then co-cultured with unmodified mesenchymal progenitors to generate compartmentalized organoids (Figure 4C). Quantitative analysis showed significantly elevated AR protein levels in AR^HI^ iPSCs compared with wild-type control (Figure 4D). Immunohistochemistry of AR^HI^ organoids confirmed robust AR and PSA expression restricted to the epithelial compartment, demonstrating successful targeting (Figure 4E). Ki67 staining further showed an increased proportion of proliferating cells in AR^HI^ organoids compared to wild-type controls, indicating that AR overexpression may enhance epithelial proliferation (Figure 4F). Transcriptomic profiles indicated elevated expression of canonical AR targets, including *KLK3*, *NKX3-1*, and *TMPRSS2*, in AR^HI^ organoids (Figure 4G). This co-culture system offers a platform that supports spatially organized prostate-like organoids and enables targeted manipulation of the epithelial lineage.

### Xeno-free human iPSC-derived prostate organoids how epithelial and stromal diversity at single-cell resolution

Single-cell RNA sequencing of week-10 wild-type organoids revealed epithelial and stromal diversity consistent with prostate lineage assembly (Figure 5A). The epithelial compartment comprised basal, luminal, club/hillock, neuroendocrine, and urethral-like populations, while the stromal compartment included smooth muscle, fibroblast, myofibroblast, pericyte, and glia-like cells.

**Figure 5.**
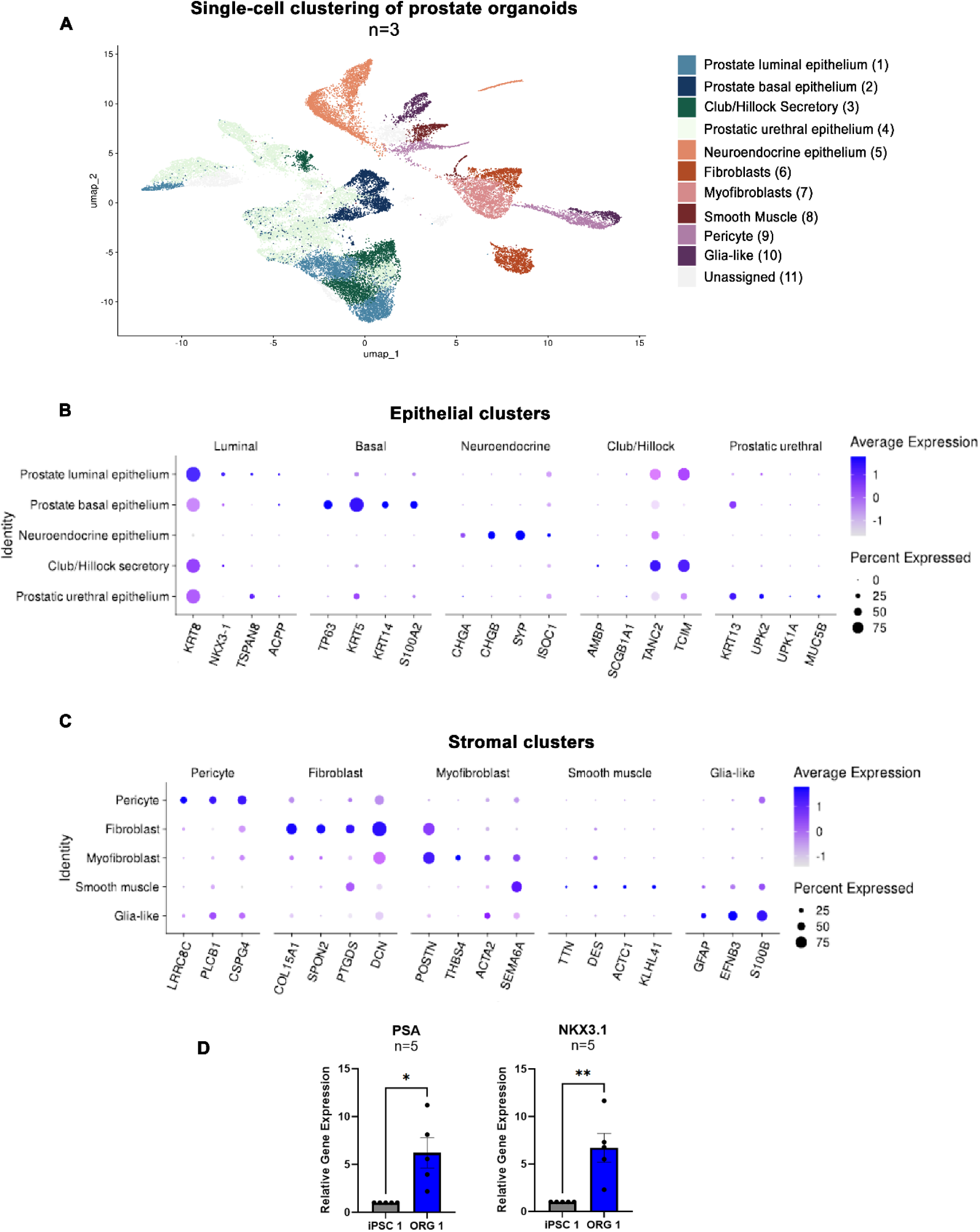
Single-cell transcriptomic profiling of WT iPSC-derived prostate organoids. (A) UMAP projection of week 10 WT organoids (*n* = 3) showing distinct epithelial and stromal clusters. Epithelial subtypes include prostate luminal, basal, club/hillock secretory, prostatic urethral, and neuroendocrine epithelium, while stromal lineages comprise fibroblasts, myofibroblasts, smooth muscle cells, pericytes, and glia-like cells. (B–C) Dot plots showing representative epithelial and stromal markers in week-10 organoids. Epithelial clusters: Luminal: *KRT8*, *NKX3-1*, *TSPAN8*, *ACPP* (androgen-responsive secretory epithelium); Basal: *TP63*, *KRT5*, *KRT14*, *S100A2* (structural/progenitor layer); Neuroendocrine: *CHGA*, *CHGB*, *SYP*, *ISOC1* (vesicle-associated hormone secretory cells resembling prostate neuroendocrine epithelium); Club/Hillock: *AMBP*, *SCGB1A1*, *TANC2*, *TCIM* (regenerative/secretory subset); Urethral-like: *KRT13*, *UPK2*, *UPK1A*, *MUC5B* (transitional epithelium of the prostatic urethra). Stromal clusters: Pericytes: *LRRC8C*, *PLCB1*, *CSPG4* (perivascular/mural support); Fibroblasts: *COL15A1*, *SPON1*, *PTGDS*, *DCN* (ECM synthesis and paracrine signaling); Myofibroblasts: *POSTN*, *THBS4*, *ACTC2*, *SEMA6A* (matrix remodeling and contractility); Smooth muscle: *TTN*, *DES*, *ACTC1*, *KLHL41* (contractile cytoskeleton); Glia-like: *GFAP*, *EFNB3*, *S100B* (neural-associated stroma). (D) qPCR (*n* = 5) showing increased *KLK3* (PSA) and *NKX3-1* in week-10 organoids (ORG1) compared with iPSCs (iPSC1); mean ± SEM; *p* < 0.05, *p* < 0.01.

Cluster annotations were validated using canonical lineage markers consistent with known prostate identities. Luminal cells expressed *KRT8*, *NKX3-1*, *TSPAN8*, and *ACPP*, whereas basal epithelial cells expressed *TP63*, *KRT5*, *KRT14*, and *S100A2*. Neuroendocrine clusters were defined by *CHGA*, *CHGB*, *SYP*, and *ISOC1*, and the club/hillock subset expressed *SCGB1A1* and *AMBP*, together with *TANC2* and *TCIM*. Urethral-like epithelial cells expressed *KRT13*, *UPK2*, and *UPK1A* (Figure 5B). Comparative analysis of major epithelial clusters against all other lineages (Supplementary Figure 4A-D) further supported their transcriptional distinction.

Stromal marker expression is shown in Figure 5C, including fibroblast genes (*COL15A1*, *SPON1*, *PTGDS*, *DCN*), myofibroblast genes (*POSTN*, *THBS4*, *ACTA2*, and *SEMA6A*), smooth muscle markers (*TTN*, *DES*, *ACTC1*, *KLHL41*), pericyte markers (*LRRC8C*, *PLCB1*, *CSPG4*), and glia-like markers (*GFAP*, *EFNB3*, *S100B*). These expression profiles correspond to expected stromal phenotypes, with myofibroblast clusters showing partial overlap between fibroblast and smooth muscle signatures, consistent with a transitional role in matrix remodeling. Orthogonal validation supported these identities. qPCR confirmed luminal differentiation by increased *KLK3* and *NKX3-1* expression (Figure 5D). Immunostaining identified desmin-positive stromal regions, *KRT13*-positive transitional epithelium, and CgA-positive neuroendocrine cells (Figure S4E). Together, these data demonstrate that xeno-free iPSC-derived prostate organoids recapitulate epithelial and stromal complexity and exhibit distinct lineage compartmentalization at single-cell resolution.

## Discussion

Organoid models have significantly advanced prostate research, providing tractable systems to study development, disease, and therapeutic responses. Tissue-derived models have enabled the expansion of benign and malignant epithelium, identification of progenitor populations, and generation of patient-derived organoids for drug testing (11,13,31,32). Despite their impact, these approaches have notable limitations, as they often lack a stromal component, are difficult to manipulate genetically after establishment, and do not easily recapitulate early developmental processes (17). Prolonged culture of primary tissue-derived organoids introduces selective pressures, and transcriptional drift affects the interpretation of results (16,33,34)

To help address these limitations, we developed a xeno-free, iPSC-based system that generates multilineage organoids with epithelial and stromal compartments. Our system captures key aspects of prostate lineage specification and tissue organization. The organoids show morphogenetic behaviors such as budding and branching, which are consistent with developmental milestones described in vivo (35).

In the monoculture system, differentiation begins with the formation of a transient mesendodermal state marked by co-expression of *SOX17* and *T (Brachyury)*. This bipotent intermediate is a recognized precursor to both mesoderm and endoderm lineages (36–38), providing a developmental foundation for generating multilineage organoids with epithelial-mesenchymal compartmentalization (39–41).

As differentiation progresses, prostate organoids derived from single iPSC lines exhibit increasing expression of key epithelial markers, including AR, NKX3-1, PSA (*KLK3*), CK8/18, and p63. Gene set enrichment analysis confirmed activation of basal and luminal transcriptional programs, in agreement with previous observations (22). Although bulk RNA-seq may not resolve rare or transitional subtypes, the presence of both basal and luminal signatures indicates successful epithelial specification. This setup is especially useful for modeling germline mutations, a growing application of iPSC-based systems in disease modeling and developmental biology (16,42).

Our observations also align with the developmental stages previously described by Cunha and Baskin (43), who showed that prostate bud initiation during the initial stages of development can occur in the absence of androgens, while branching and epithelial maturation require androgen signaling. Consistently, we found that organoids cultured without dihydrotestosterone (DHT) exhibited reduced expression of epithelial markers such as CK8/18, p63, and PSA, while stromal markers like α-SMA and vimentin remained robust or elevated. This suggests that stromal differentiation may be less dependent on androgen signaling than epithelial development.

To enable lineage-specific manipulation, we developed a co-culture system in which epithelial and stromal compartments are specified independently. Genetic modification can be introduced prior to differentiation into definitive endoderm (*FOXA2⁺*, *SOX17⁺*), which is subsequently combined with wild-type mesenchymal progenitors (*NCAM⁺*). This approach enables targeted gene editing within the epithelial lineage while preserving a physiologically relevant stromal environment. The resulting organoids displayed distinct epithelial (*PSA⁺*, *AR⁺*) and stromal (*VIM⁺*, *α-SMA⁺*) domains, consistent with organized prostate-like architecture. To demonstrate the utility of this system for disease modeling, an *AR* transgene was introduced into iPSCs prior to differentiation. The resulting organoids showed robust AR and PSA expression restricted to the epithelial compartment, while stromal cells remained unmodified. Increased Ki67 staining in AR^HI^ organoids is consistent with increased epithelial proliferation, a feature relevant to both normal prostate development and *AR*-driven tumorigenesis. Transcriptomic profiling revealed higher read counts for canonical *AR* target genes (*KLK3*, *NKX3-1*, *TMPRSS2*) in AR^HI^ organoids. Collectively, these findings demonstrate that the co-culture system enables controlled, lineage-specific modeling of oncogenic drivers within a spatially organized epithelial–stromal environment.

Single-cell RNA sequencing of wild-type organoids revealed basal (*TP63*, *KRT5*, *KRT14*), luminal (*NKX3-1*, *TSPAN8*, *ACPP*), and neuroendocrine (*CHGA*, *CHGB*, *SYP*) populations, together with club/hillock clusters representing secretory epithelial subsets. These populations collectively reflect the epithelial diversity of the developing prostate. As expected for probe-based single-cell approaches, transcripts with low abundance or unstable 3′ ends, such as *KLK3* (encoding PSA), showed reduced detection despite clear luminal identity confirmed by qPCR and immunostaining (44,45). The coexistence of progenitor and differentiated epithelial phenotypes suggests an actively remodeling epithelium, and the balance between these states can be modulated by differentiation timing or inductive cues. This dynamic organization provides a physiologically relevant framework to investigate developmental reactivation processes linked to prostate cancer.

Stromal identities were supported by single-cell markers for smooth muscle (*ACTA1*, *DES)*, fibroblast (*DCN*, *COL15A1*), pericyte (*PLCB1*, *CSPG4*), and glia-like populations (*S100B*, *EFNB3*), with clear module enrichment across stromal clusters. Desmin immunostaining further corroborated smooth-muscle differentiation in the organoid stroma (46), aligning the transcriptomic calls with spatial protein expression. Although stromal subtypes partially overlapped in marker usage, their gene modules were distinct and consistent across replicates, supporting the robustness of lineage representation.

The single-cell landscape captured a fuller breadth of prostate lineage programs, including the prostatic urethra, that share a common embryonic development from the urogenital sinus epithelium (35,47,48). Specifically, there is a subset of epithelial cells expressing *KRT13*, *UPK2*, and *UPK1A* is consistent with the cells of the prostatic urethra and proximal ducts, as demonstrated by recent reports of whole human prostate single-cell characterization (25,49,50). The interaction between these cells, along with club and hillock epithelial cell types, provide a model to explore their potential role in BPH (25).

This iPSC-derived system complements tissue-derived models by providing a reproducible, developmentally informed platform with matched epithelial and stromal organization. Recent Matrigel-free patient-derived organoid studies have shown that eliminating animal-derived matrices preserves prostate cancer lineage fidelity and reduces transcriptional artefacts (51), underscoring the importance of defined, xeno-free conditions for translational relevance. Because the organoids generated here include transitional states and a urethral-like program, they provide a useful framework to investigate processes relevant to benign prostatic hyperplasia (BPH), including the reactivation of developmental pathways (52,53). The modular design, which allows lineage-specific genetic manipulation before aggregation, further enables mechanistic studies of how defined drivers and cues shape fate choices in a human system and support studies on differentiation, lineage specification, BPH, and malignant transformation.

### Limitations of the study

This report presents a defined protocol and an initial experimental series spanning two differentiation time points, weeks 6 and 10. Probe-based single-cell RNA sequencing was conducted at week 10 to characterize lineage diversity once epithelial and stromal compartments had become clearly distinguishable, providing a representative snapshot of prostate-like tissue organization rather than a full developmental trajectory. Earlier changes were assessed by bulk transcriptomics and immunostaining, which complement the single-cell dataset. Cell composition reflects the patterning cues and culture duration used here and can be further refined in future experiments to favor specific epithelial outcomes. The current system does not yet include immune or vascular components, so those interactions are not yet represented.

## STAR Methods

### Patient material

All surgical specimens were collected in accordance with local ethical and regulatory guidelines, with written informed consent from patients (Newcastle REC 2003/11 and Human Tissue Authority License 12,534, Freeman Hospital, Newcastle upon Tyne, United Kingdom). The anonymized samples were labelled iPSC 1, iPSC 2, and iPSC 3, corresponding to patients aged 82, 81, and 67 years at the time of surgery.

### iPSC Generation

Enriched cultures of 1 × 10⁵ prostate stromal cells were seeded in 12-well plates and transduced using the CytoTune™-iPS 2.0 Reprogramming Kit (Thermo Fisher Scientific, A16517) according to the manufacturer’s instructions. Induced pluripotent stem cell (iPSC) colonies were first established on an inactivated primary mouse embryonic fibroblast feeder layer, then transitioned to a feeder-free system as described below.

### iPSC Culture

Human iPSCs were cultured in six-well plates coated with Vitronectin (Thermo Fisher Scientific, A14700) using mTeSR™1 medium (StemCell Technologies, 85850). The culture medium was replaced daily. Cells were allowed to grow for 4–5 days, before either passaging or initiating differentiation. For passaging, cells were treated with Versene solution (Thermo Fisher Scientific, 15040066) at 37°C for 3–5 minutes, and then transferred to fresh Vitronectin-coated plates at a ratio of 1:3 to 1:6. All cultures were maintained in a humidified environment at 37°C with 5% CO2.

### iPSC Characterization

Pluripotency markers in iPSC colonies were detected using immunocytochemistry and flow cytometry. Colonies were fixed in 4% paraformaldehyde (Sigma-Aldrich, 47608), permeabilized with 0.25% Triton X-100 (Sigma-Aldrich, T8787), and blocked with 10% FBS and 1% bovine serum albumin (Sigma-Aldrich, A3311) before staining with anti-human SOX2 (Merck Millipore, MAB4343, 1:200) and anti-human OCT4 (Merck Millipore, MABD76, 1:200). Secondary staining was performed using Alexa Fluor 647 goat anti-mouse (Thermo Fisher Scientific, A21235, 1:400), followed by DAPI counterstaining at 1 µg/ml (Thermo Fisher Scientific, D1306), and imaged with a Leica DM6 microscope. For flow cytometry, iPSCs were dissociated with Accutase (StemCell Technologies, 07920), stained with TRA-1-60-FITC (Merck Millipore, FCMAB115F, 1:60) and NANOG-Alexa Fluor® 647 (Cell Signaling Technology, 5448S, 1:150), and analyzed using a Fortessa X20 system. For in vitro three-germ-layer differentiation, iPSCs were treated with Dispase (StemCell Technologies, 07923), cultured in differentiation media containing DMEM-F12 (Thermo Fisher Scientific, 11330), 20% FBS (Thermo Fisher Scientific, 10270), and stained for germ-layer markers using the antibodies AFP (Thermo Fisher Scientific, MA5-14666), TUBB3 (BioLegend, 801201), and α-SMA (Abcam, ab5694). Negative controls used only secondary antibodies.

### qPCR

Reverse transcription was performed on 1µg of RNA using MMLV reverse transcriptase (Promega, M1701) following the manufacturer’s instructions. Quantitative PCR was performed using the PowerTrack Master Mix (ThermoFisher, A46109) according to manufacturer’s guidance. Expression of markers (Supplementary Table 1) was normalized to *RPL13A*.

### iPSC Engineering

The full-length androgen receptor (ARFL) cDNA was cloned into the MYC_pLX307 vector (Addgene, 98363) using T4 ligase (Thermo Fisher Scientific, IVGN2104), after removing the MYC sequence with the restriction enzymes NheI and SpeI (New England Biolabs, R3131S and R3133S). Bacterial transformation was carried out in Stbl3™ chemically competent *E. coli* (Thermo Fisher Scientific, C737303), and plasmid purification was performed using the Invitrogen miniprep kit (Thermo Fisher Scientific, K210011). The presence of ARFL in the plasmid was confirmed through Sanger sequencing and diagnostic restriction digestion. The ARFL_pLX307 plasmid was packaged into lentivirus and resuspended in culture media following ultracentrifugation. iPSCs were transduced one day post-passage with 250 µl of viral particle solution for 48 hours. Cells carrying the plasmid were selected using blasticidin (1 µg/ml), followed by clonal selection to ensure that all cells contained the AR construct.

## Organoid Generation

### Monoculture

#### DE Spheroid Formation (days 0 to 3)

Cells were first washed with PBS and incubated with 1 mL of Accutase Reagent (StemCell Technologies, 07920) for 3 minutes at 37°C to facilitate detachment. After incubation, 1 mL of mTeSR medium (StemCell Technologies, 85850) was added, and the cells were gently pipetted 1–3 times to generate a single-cell suspension. The suspension was transferred to a 15 mL conical tube and centrifuged at 300 × g for 5 minutes. Following centrifugation, the supernatant was discarded, and cells were resuspended in 1 mL of mTeSR medium per well collected. Viable cell concentration was determined using Trypan Blue (Thermo Fisher Scientific, 15250061) and a hemocytometer. The suspension was then adjusted to 1,000 cells per well in 100 μL of mTeSR medium supplemented with 10 μM ROCK inhibitor (StemCell Technologies, 72304). Cells were plated in ultra-low attachment 96-well plates (Corning Costar, 7007) and immediately centrifuged at 250 × g for 10 minutes at 20°C to encourage uniform cell settling. Plates were incubated at 37°C with 5% CO₂ and 95% humidity. After 24 hours, daily media changes were performed for 3 days using 100 μL of differentiation medium consisting of RPMI 1640 (Thermo Fisher Scientific, 11875093) supplemented with 1% Penicillin-Streptomycin (Thermo Fisher Scientific, 15140122), 1% GlutaMAX (Thermo Fisher Scientific, 35050061), 100 ng/mL Activin A (StemCell Technologies, 78034), 1× B-27, and 10 nM DHT.

#### Prostate Organoid Specification (Days 4–8)

From days 4 to 8, prostate specification treatment was initiated by adding 100 μL of the following medium to each well daily: RPMI 1640 (Thermo Fisher Scientific, 11875093) supplemented with 1% Pen-Strep (Thermo Fisher Scientific, 15140122), 1% GlutaMAX (Thermo Fisher Scientific, 35050061), 50 ng/mL RSPO1 (Thermo Fisher Scientific, A42586), 10 ng/mL EGF (Thermo Fisher Scientific, PHG0315), 10 ng/mL FGF10 (Thermo Fisher Scientific, 100-26), 1x B27 (Thermo Fisher Scientific, 17504044), and 10 nM DHT.

#### Prostate Organoid Differentiation (Day 9 onward)

Starting from day 9, prostate organoid differentiation treatment was initiated, with 100 μL per well added every other day. The treatment medium consisted of Advanced DMEM (Thermo Fisher Scientific, 12491015) with 1% Pen-Strep and 1% GlutaMAX, 50 ng/mL RSPO1, 10 ng/mL EGF, 10 ng/mL FGF10, 5 ng/mL BMP4 (Thermo Fisher Scientific, PHC9534), 5 ng/mL IGF1 (Thermo Fisher Scientific, 78142), 10 mM HEPES (Thermo Fisher Scientific, 15630056), 1X B27, 10 nM DHT, and 1 μL/mL Amphotericin B (Thermo Fisher Scientific, 15290018). Organoids were harvested at weeks 6 and 10 for further analysis.

### Co-culture

iPSC-derived definitive endoderm cells and mesenchymal progenitors were dissociated using Accutase (StemCell Technologies, 07920), centrifuged at 300 × g, and counted using Trypan Blue (Thermo Fisher Scientific, 15250061). The two cell types were combined at a 3:1 ratio (mesenchymal: endodermal) to a final density of 1,000 cells per well in 100 μL of media (as described in Step 2 of the monoculture protocol). Cells were seeded into ultra-low attachment 96-well plates, (Corning Costar, 7007) and immediately centrifuged at 250 × g for 10 minutes to encourage aggregation.

Definitive endoderm: iPSCs were dissociated into single cells using Gentle Cell Dissociation Reagent (StemCell Technologies, 07174) following the manufacturer’s instructions. A total of 5 × 10⁵ cells were seeded into Vitronectin-coated 6-well plates in mTeSR1 medium, (StemCell Technologies, 85850) supplemented with 10 μM ROCK inhibitor (StemCell Technologies, 72304). After 24 hours, the medium was replaced with RPMI 1640 (Thermo Fisher Scientific, 11875093) containing 100 ng/mL human recombinant Activin A (StemCell Technologies, 78034) and 2.5 μM CHIR99021 (Tocris, 4423), without additional supplements. On Day 2, cells were washed once with PBS and cultured in RPMI 1640 supplemented with 100 ng/mL Activin A and B-27 minus insulin (Thermo Fisher Scientific, A1895601; 1:50 dilution). On Day 3, the medium was changed to RPMI 1640 containing 100 ng/mL Activin A and B-27 complete (Thermo Fisher Scientific, 17504044; 1:50 dilution). Definitive endoderm cells were harvested after 72 hours for downstream use.

Mesenchymal progenitors: iPSCs were differentiated into mesenchymal progenitors over 21 days using the STEMdiff™ Mesenchymal Progenitor Kit (StemCell Technologies, 05240), following the manufacturer’s instructions. The protocol included three key stages: passaging iPSCs, induction of early mesodermal progenitors, and maturation into mesenchymal progenitor cells. Each stage involved coating culture plates, preparing single-cell suspensions, performing medium changes, and monitoring confluency and viability using Trypan Blue (Thermo Fisher Scientific, 15250061).

## Real-time morphological analysis

Plates were placed in the Incucyte® Live-Cell Analysis System and scanned at 4X magnification, capturing one image per well. The phase contrast/Brightfield channels were selected, with a spheroid growth scan type and a scan interval of every 6 hours. Morphological changes were counted independently on days 3, 7, 14, 21, 28, 35, and 42. This assessment focused on two key features: the number of buds, defined as round structures emerging from the organoids, and branching, which refers to the expansion of buds or the connection of multiple buds together.

## Histological analysis

### Fixation, Processing, and Paraffin Embedding of Organoids

Organoids were harvested at weeks 6 or 10, washed in PBS, and fixed in 10% neutral buffered formalin (NBF) for 1 hour at room temperature. Following fixation, samples were washed again with PBS, and Histogel (Thermo Fisher Scientific or equivalent) was melted in Eppendorf tubes using a dry heat block at 100°C for 5 minutes, then mixed with organoids and transferred into embedding molds. Molds were left at 4°C overnight to solidify. The following day, solidified Histogel plugs were transferred into tissue cassettes and placed in 70% ethanol for storage or immediate processing. Samples were processed through a graded alcohol and xylene series using a Leica tissue processor, then embedded in paraffin using a standard embedding station. Paraffin blocks were sectioned at 4– 5 μm thickness using a microtome, and sections were mounted onto charged glass slides for downstream histological or immunohistochemical analysis.

### Immunohistochemistry Protocol for FFPE Tissue Slides

Formalin-fixed, paraffin-embedded (FFPE) tissue slides were deparaffinized in xylene and rehydrated through a graded ethanol series. Antigen retrieval was performed using heat-induced epitope retrieval in citrate buffer with a pressure-based decloaking chamber, heated to 121°C for 20 minutes. Slides were then cooled and rinsed under running water. Endogenous peroxidase activity was blocked using 3% hydrogen peroxide, and nonspecific binding was minimized by incubating the slides with blocking serum. Primary antibodies (listed in Supplementary Table 2) were applied and incubated, followed by horseradish peroxidase (HRP)-conjugated secondary antibodies. Detection was achieved using DAB staining. Slides were counterstained with hematoxylin, dehydrated through ethanol, cleared in xylene, and mounted with DPX. Slides were scanned using the Aperio system for digital image capture and analysis.

## Gene expression analysis

### Bulk RNA-seq Analysis

RNA libraries were prepared using the NEBNext Ultra II RNA Library Prep Kit for Illumina (New England Biolabs) and sequenced on an Illumina NovaSeq platform (2×150 bp paired end), generating approximately 20 million reads per sample. Raw data quality was assessed using FASTQC v0.12.1 (54), and adapter trimming was performed using Trim Galore v0.6.10 (55). Transcript quantification was carried out using Salmon v1.10.1 (56), with GENCODE Release 46 (GRCh38.p14) as the reference and the GRCh38 primary genome assembly used as a decoy to improve mapping accuracy. Transcript-level estimates were imported into R v4.3.2 using tximport v1.30.0 (57) from Bioconductor v3.18 (58) and summarized to gene-level counts. Transcript and gene identifiers were based on Ensembl release 111 (59).

Differential gene expression analysis was performed using DESeq2 v1.42.1 (60), with genes considered significant at adjusted p-value (padj) < 0.01 and absolute log2 fold change > 1. Principal Component Analysis (PCA) was conducted on variance-stabilized transformed (VST) data using the top 500 most variable genes. Volcano plots were generated from DESeq2 output to visualize the magnitude and significance of differential expression.

Gene Ontology (GO) enrichment was performed on differentially expressed genes (DEGs) using clusterProfiler v4.10.1 (61), targeting Biological Process (GO BP) terms with p-value < 0.05 and q-value < 0.2 (Benjamini-Hochberg correction). Gene Set Enrichment Analysis (GSEA) was also conducted on ranked DEGs using prostate-relevant hallmark, curated, and ontology gene sets from msigdbr v7.5.1 (62), with adjusted p-value < 0.05.

Bulk RNA-seq datasets of sorted basal and luminal prostate cells from primary prostate epithelial tissue (63) were used to derive reference signatures. The top 50 DEGs (padj < 0.01) were selected to create curated gene sets. These were used in GSEA to assess basal and luminal enrichment across organoid samples.

For visualization, VST-normalized expression of epithelial, stromal, neuroendocrine, and extracellular matrix markers was averaged across replicates, compiled into matrices, and plotted as heatmaps using the pheatmap R package v1.0.12 (64).

### Single-cell RNA-seq Analysis

Single-cell libraries were generated using the 10x Genomics Fixed RNA Profiling v1 chemistry with the Human Transcriptome Probe Set v1.0.1, with sequencing performed by the Genomics Core Facility, Newcastle University. Reads were aligned to the GRCh38-2020-A reference transcriptome using Cell Ranger v8.0.0 (65) processed by the Bioinformatics Support Unit.

Gene expression matrices were imported into Seurat v4.4.0 (66) for downstream analysis. Cells were filtered to exclude those with fewer than 600 UMIs, fewer than 400 detected genes, a gene/UMI log10 ratio below 0.80, or mitochondrial content exceeding 10%. Genes expressed in fewer than 200 cells were excluded. Visualizations including histograms, violin plots, and scatter plots were used to guide quality control.

Each replicate was independently processed through the standard Seurat pipeline, including normalization, variable feature identification, scaling, and clustering. Doublets were detected and removed using DoubletFinder v2.0.4 (67). After quality filtering, replicate Seurat objects were merged, and cell cycle effects were regressed out using the default gene sets in Seurat.

Dimensionality reduction was performed using Principal Component Analysis (PCA), followed by nearest neighbor identification and clustering. UMAP was used for visualization. Clusters were manually annotated using known prostate cell markers, and top markers within each cluster were identified based on a log2 fold change threshold of 0.25. The top 25 genes per cluster were used to refine annotations.

## Acknowledgements

This work was primarily supported by the National Centre for the Replacement, Refinement and Reduction of Animals in Research (NC3Rs; project NC/W002396/1, Training Fellowship awarded to Adriana Buskin). We also acknowledge additional support from Prostate Cancer UK (project TLD-CAF23-007) and the Prostate Cancer Foundation (project 18CHAL11). Acquisition of the Live-Cell Analysis System was enabled by an MRC equipment grant with the support of Dr. Kelly Coffey. Financial support to Majlinda Lako from the IMI2 StemBANCC initiative is also gratefully acknowledged.

## Author Contributions

Conceptualization: A.B., R.H.; Methodology: A.B.; Investigation: L.J.W., L.W., R.Ho., S.S., A.C.H., R.N., R.Hu., J.C., A.B.; Formal analysis: N.S., M.T., S.B., A.B.; Data curation: N.S., M.T., S.B.; Visualization: N.S., M.T., A.B.; Resources: A.C.H., L.G., M.L.; Supervision: E.S., C.R., A.B.; Funding acquisition: A.B.; Writing: original draft: A.B.; Writing: review & editing: N.S., M.T., Q.L., B.S., S.H., D.S., R.H., A.B.

---

**Supplementary Figure 1.**
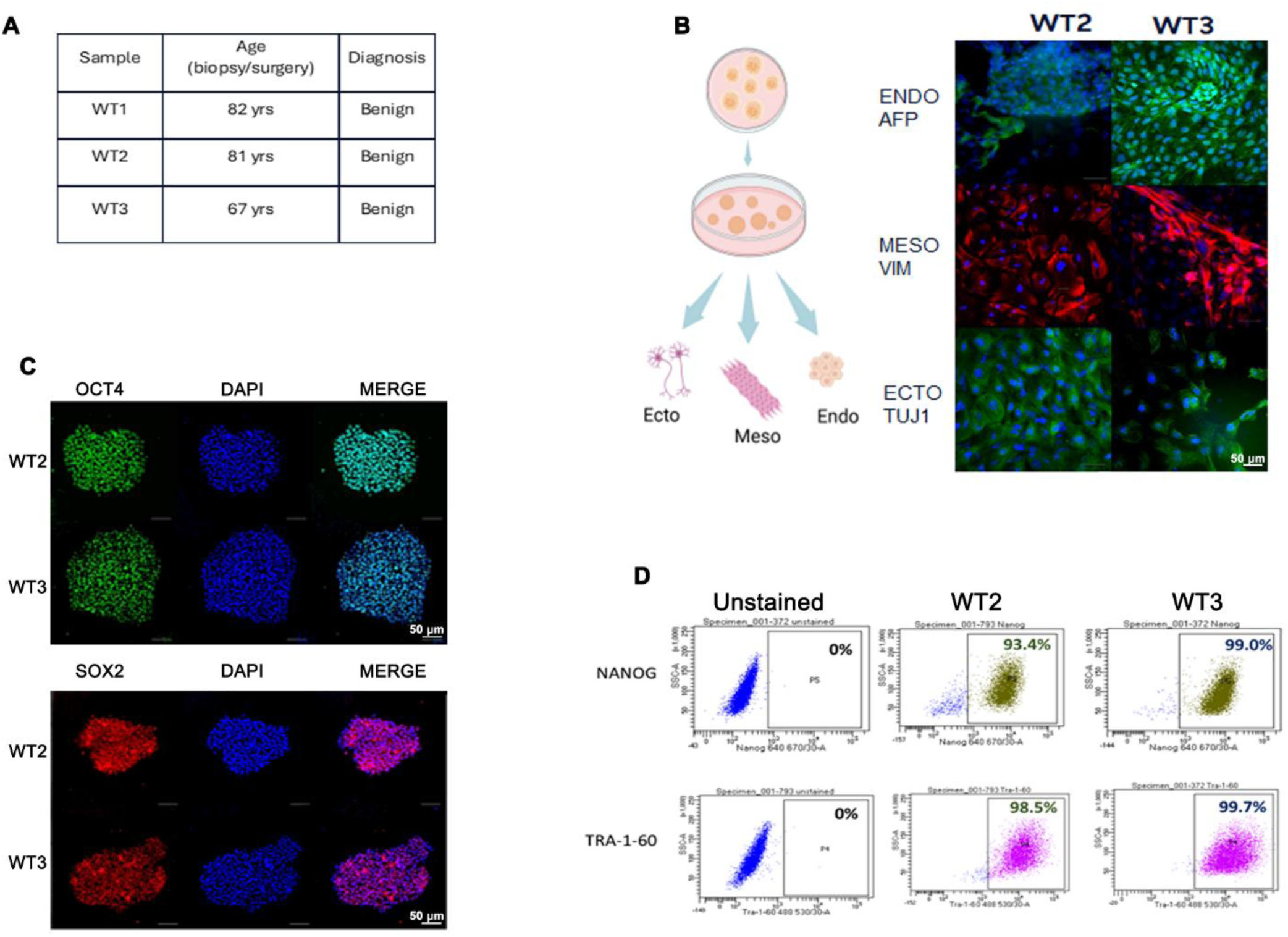
Generation and characterization of iPSCs used for prostate differentiation. (A) Overview of the three iPSC lines used in this study. WT1 data were previously published in Hepburn et al. (22). (B) Trilineage differentiation assay demonstrating that iPSCs can generate endoderm, mesoderm, and ectoderm derivatives. (C) Immunofluorescence staining showing expression of pluripotency markers OCT4 and SOX2. (D) Flow cytometry analysis confirming expression of NANOG and TRA-1-60, supporting the pluripotent status of the iPSCs.

**Supplementary Figure 2.**
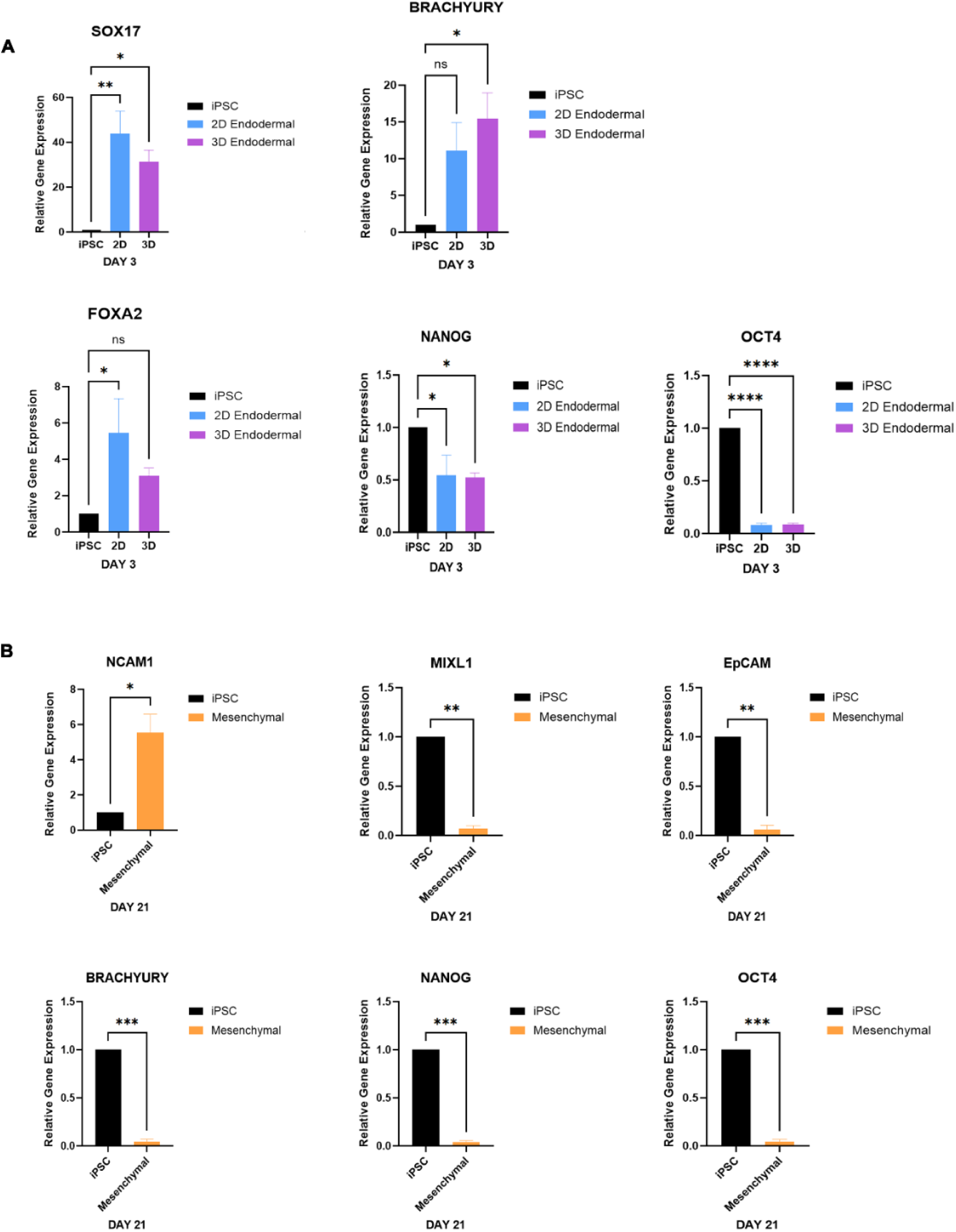
Validation of endodermal and mesenchymal progenitor differentiation from iPSCs. (A) Expression of markers *SOX17*, *T* (Brachyury), *FOXA2*, *NANOG* and *OCT4* in 3D and 2D cultures via qPCR. Expression of *SOX17* and *T* (Brachyury) in 3D spheroids suggests the presence of a mesendodermal intermediate. Downregulation of *NANOG* and *OCT4* indicates loss of iPSC identity after differentiation. n = 4 biological replicates per condition. (B) By day 21, differentiation to mesenchymal progenitors is supported by upregulation of *NCAM1* and loss of epithelial and pluripotency markers, with reduced *EPCAM*, *NANOG*, *OCT4*, and *T* (Brachyury). *MIXL1*, a transient early mesoderm marker, is downregulated, consistent with progression beyond early mesoderm to a committed mesenchymal fate. n = 3 biological replicates per condition. Bars represent mean ± SD of relative gene expression quantified by 2^−ΔΔCt, normalized to *RPL13A* and calibrated to iPSC = 1. Pairwise comparisons were performed using two-sided Welch *t*-tests. Significance: ns p ≥ 0.05, * p < 0.05, ** p < 0.01, *** p < 0.001, **** p < 0.0001.

**Supplementary Figure 3.**
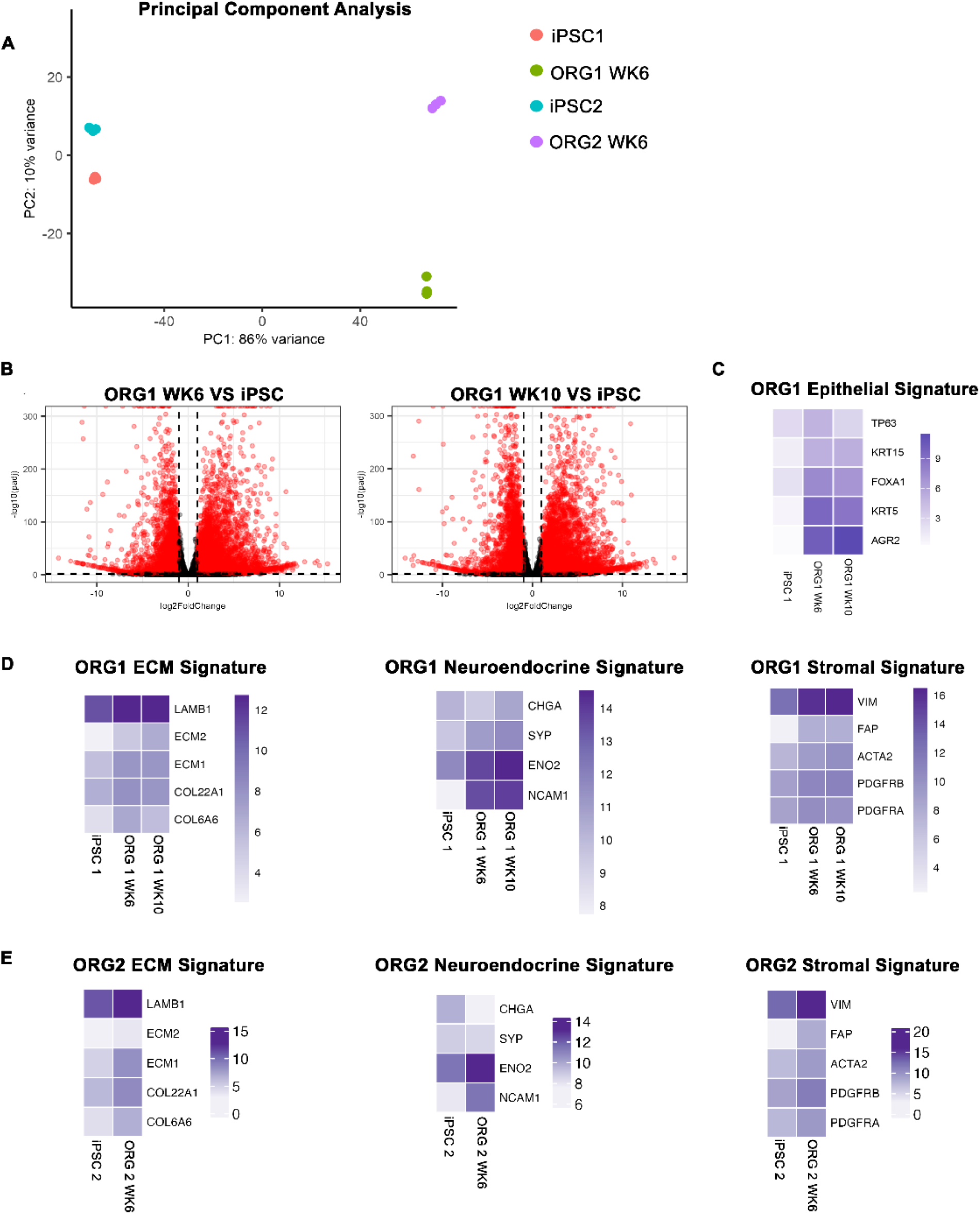
Bulk RNA-seq reveals transcriptional changes and pathway enrichment associated with prostate organoid differentiation. (A) Principal component analysis (PCA) shows transcriptional divergence between undifferentiated iPSCs and organoids at week 6, indicating distinct gene expression profiles following differentiation. (B) Volcano plots showing differentially expressed genes in organoids (ORG1) at weeks 6 and 10 compared with iPSCs, highlighting dynamic transcriptional changes during differentiation. (C) Heatmap of epithelial gene signatures in ORG1 organoids, week 6 and week 10 timepoints. (D) Heatmaps of selected gene signatures in ORG1 organoids, including extracellular matrix (ECM) genes (*LAMB1*, *ECM1*, *ECM2*, *COL22A1*, *COL6A6*), neuroendocrine markers (*CHGA*, *SYP*, *ENO2*, *NCAM1*), and stromal genes (*VIM*, *FAP*, *ACTA2*, *PDGFRB*, *PDGFRA*) across iPSC, week 6, and week 10 timepoints. (E) Equivalent ECM, neuroendocrine, and stromal signatures observed in ORG2 organoids at week 6 support reproducibility of lineage specification across biological replicates.

**Supplementary Figure 4.**
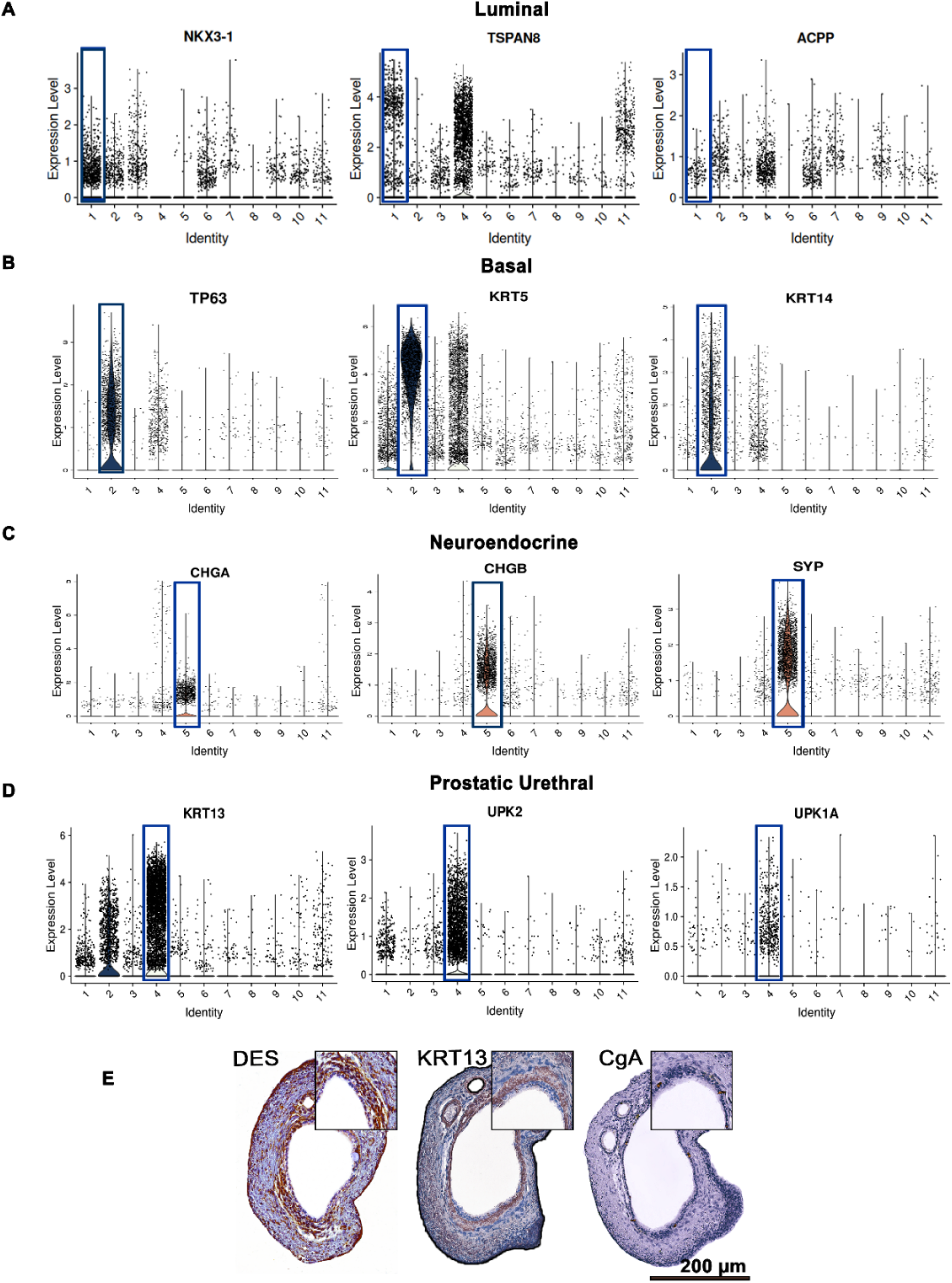
Validation of prostate lineage acquisition in WT iPSC-derived organoids. (A) Violin plots of luminal markers *NKX3-1*, *TSPAN8*, and *ACPP*, showing expression in the luminal cluster and detectable levels in other epithelial populations. (B) Violin plots of basal markers *TP63*, *KRT5*, and *KRT14*, showing expression in the basal cluster. (C) Violin plots of neuroendocrine markers *CHGA*, *CHGB*, and *SYP*, showing expression in the neuroendocrine cluster. (D) Violin plots of prostatic urethral cell markers *KRT13, UPK2*, and *UPK1A*, showing expression in the prostatic urethral epithelium cluster. (E) Immunohistochemistry for desmin (stromal), keratin 13 (urethral epithelium), and chromogranin A (neuroendocrine) showing spatial organization of prostate-relevant lineages. Scale bar, 200 µm. **Cluster identities:** (1) Prostate luminal epithelium, (2) Prostate basal epithelium, (3) Club/Hillock secretory, (4) Prostatic urethral epithelium, (5) Neuroendocrine epithelium, (6) Fibroblast, (7) Myofibroblast, (8) Smooth muscle, (9) Pericyte, (10) Glia-like, (11) Unassigned.

**Table 1.**
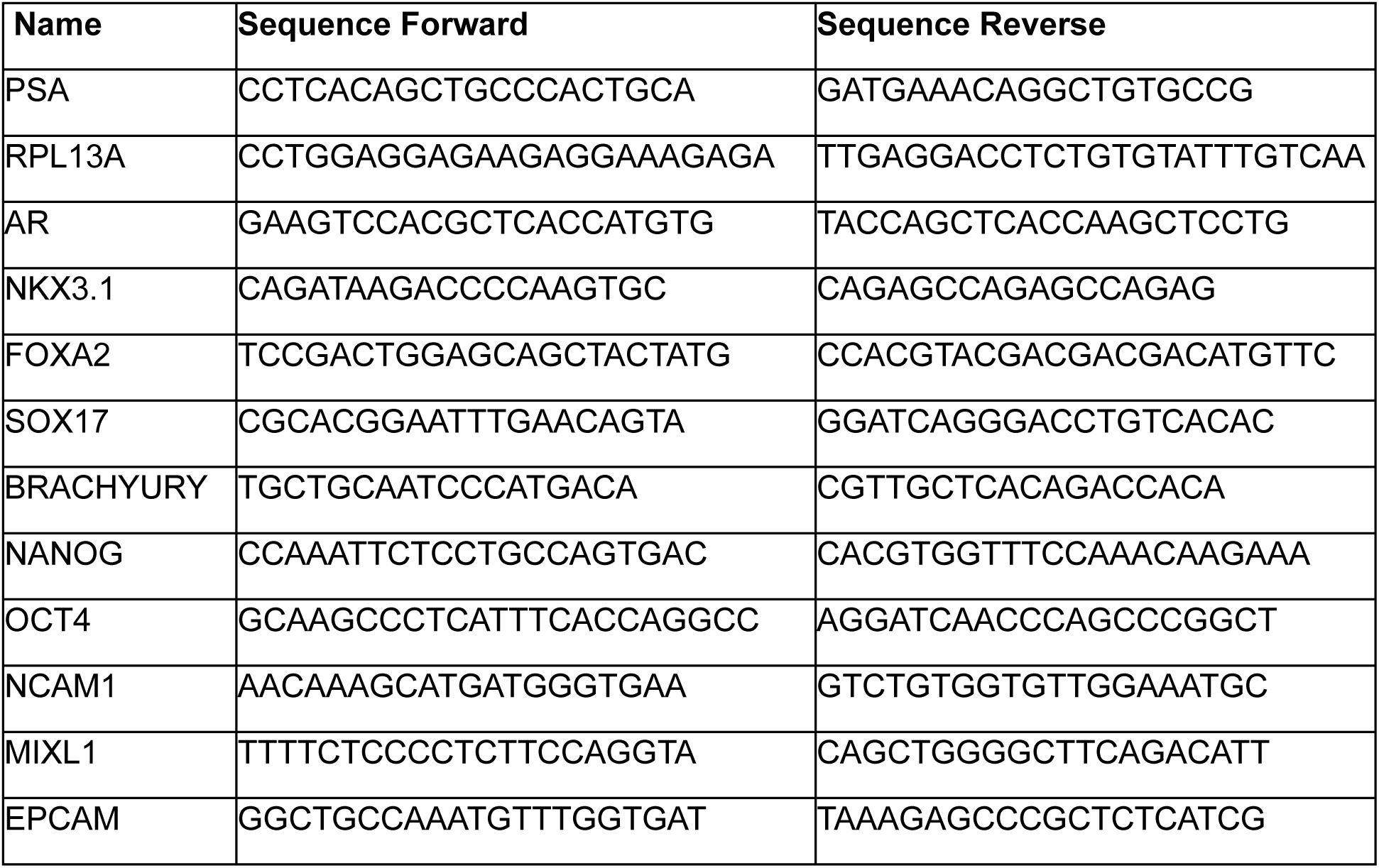
Primer sequences used in this study. Related to STAR methods.

**Table 2.** Antibodies used in this study. Related to STAR methods.

| Antibody | Dilution | Manufacturer |
| --- | --- | --- |
| AR | 1:50 | BD Biosciences 554225 (Clone AR441) |
| NKX3-1 | 1:300 | Origene UltraMAB UM800088 |
| CK8-18 | 1:300 | Abcam ab17139 |
| P63 | 1:50 | Leica Novocastra NCL-L-p63 |
| Vimentin | 1:500 | Cell Signaling D21H3 |
| α-SMA | 1:50 | Abcam ab7817 |
| PSA | 1:50 | Santa Cruz SC-7316 |
| CgA | 1:200 | Abcam ab5283 |
| Desmin | 1:2000 | Proteintech 16520-1-AP |
| KRT13 | 1:100 | Proteintech 10164-2-AP |

